# Detection of task-relevant and task-irrelevant motion sequences: application to motor adaptation in goal-directed and whole-body movements

**DOI:** 10.1101/339648

**Authors:** Daisuke Furuki, Ken Takiyama

## Abstract

Motor variability is inevitable in our body movements and is discussed from several various perspectives in motor neuroscience and biomechanics; it can originate from the variability of neural activities, it can reflect a large degree of freedom inherent in our body movements, it can decrease muscle fatigue, or it can facilitate motor learning. How to evaluate motor variability is thus a fundamental question in motor neuroscience and biomechanics. Previous methods have quantified (at least) two striking features of motor variability; the smaller variability in the task-relevant dimension than in the task-irrelevant dimension and the low-dimensional structure that is often referred to as synergy or principal component. However, those previous methods were not only unsuitable for quantifying those features simultaneously but also applicable in some limited conditions (e.g., a method cannot consider motion sequence, and another method cannot consider how each motion is relevant to performance). Here, we propose a flexible and straightforward machine learning technique that can quantify task-relevant variability, task-irrelevant variability, and the relevance of each principal component to task performance while considering the motion sequence and the relevance of each motion sequence to task performance in a data-driven manner. We validate our method by constructing a novel experimental setting to investigate goal-directed and whole-body movements. Furthermore, our setting enables the induction of motor adaptation by using perturbation and evaluating the modulation of task-relevant and task-irrelevant variabilities through motor adaptation. Our method enables the identification of a novel property of motor variability; the modulation of those variabilities differs depending on the perturbation schedule. Although a gradually imposed perturbation does not increase both task-relevant and task-irrelevant variabilities, a constant perturbation increases task-relevant variability.

## Introduction

In our daily life, we can repeatedly achieve desired movements, such as grasping a cup, throwing a ball, and playing the piano. To achieve the desired movements, our motor system needs to resolve at least two difficulties inherent in our body motion [1]. A difficulty is movement variability. Due to the variability inherent in various stages such as sensing sensory information [2], neural activities in motor planning [3], or muscle activities in motor execution [4], even sophisticated athletes and musicians cannot repeat the same movements. Our motor systems somehow tame those variabilities to achieve the desired movements [5]. Another difficulty is a large degree of freedom (DoF) inherent in our motor system [1, 6]. The number of joints, muscles, and neurons are more than necessary to achieve the desired movements, resulting in an infinite number of joint configurations, muscle activities, and neural activities that can correspond to the desired movement [7-10]. Our motor systems somehow resolve those difficulties (i.e., variability and a large DoF) and generate the desired movements.

Although it remains unclear how we tame movement variability, a possible answer is the decomposition of motor variability into task-relevant and task-irrelevant variabilities. We compensate for a portion of motor variability that is relevant to achieve the desired movements (i.e., task-relevant variability) [11-15] - simultaneously, we do not significantly compensate for a portion of the variability that is irrelevant to achieve the desired movements (i.e., task-irrelevant variability). The compensation of task-relevant variability can be observed in movement kinematics [11-14], muscle activities [16, 17], and neural activities [15]. This striking feature of our motor variability enables the achievement of the desired movements in the presence of movement variability.

Several studies have developed techniques to evaluate task-relevant and task-irrelevant variabilities. The uncontrolled manifold (UCM) evaluates the task-relevant and task-irrelevant variabilities (mainly) in joint angles and angular velocities. The method focuses on kinematic parameters relevant to task achievements, such as hip joint position in stand-and-sit motion [11] or hand position in arm-reaching movements [18]. The Jacobian matrix, the derivatives of those kinematic parameters concerning joint angles or angular velocities, enables the definition of the null space around the joint angles or angular velocities averaged across trials. The variability along the null space can be defined as the task-irrelevant variability. Those previous studies have revealed that the task-relevant variability of joint angles and angular velocities is less than the task-irrelevant variability. Notably, the UCM focused on forward kinematics that map joint angles and angular velocities into joint positions and velocities in the external coordinate. In contrast, tolerance, noise, and covariation analysis (TNC) [13] and goal-equivalent manifold analysis (GEM) [14] focused on task functions that define the relationship between kinematic parameters and task performance. For example, a thrown dart or ball can be modeled as a parabola. When the release position and velocity in the vertical axis are *p* and *v*, respectively, the maximum height of the released dart or ball can be written as 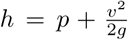, where *g* is the gravitational acceleration. When controlling *h* is a task, the relation among *h*, *p*, and *v* can be task function. For example, with a small value *d* (i.e., *d*^2^ *≃* 0), a slight change in the release position 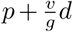 and the release velocity *v - d* does not cause any change in *h*; thus, the variability along these slight changes to be regarded as the task-irrelevant variability. TNC and GEM evaluate the task-relevant and task-irrelevant variabilities based on the task functions.

Those techniques have pros and cons. UCM enables the evaluation of motion sequence, but it does not consider the task function. The framework is thus not always suitable for the situation when kinematic parameters are nonlinearly relevant to task achievements, such as the quadratic function of *v* in the parabolic mentioned above. Because forward kinematics are nonlinear functions of joint angles and angular velocities, UCM requires local linear approximation around representative joint angles or angular velocities based on the Jacobian matrix. Due to the linear approximation, UCM assumed the variability around the kinematics averaged across all the trials. The approximation results in difficulty simultaneously considering the motor variability when the averaged kinematics change, such as those before, during, and after motor learning (although it is possible to discuss those situations separately [18]). GEM also considers the local linear approximation of the nonlinear task function; thus, the method also considers the variability around the task parameters averaged across all the trials. Although GEM can deal with the task function, it is difficult to consider motion sequence in several cases. For example, to consider the motion sequence, the GEM framework needs to define how the dart or ball position and velocity 100 msec before the release are relevant to the maximum height in the example, as mentioned above. TNC enables the simultaneous discussion of the motor variability before, during, and after motor learning because the framework captures the whole variability in a nonparametric manner without local linear approximation; however, TNC is not always suitable for considering motion sequence based on a similar reason to GEM - it requires an explicit definition of the task function. In total, each method has pros and cons; thus, no single framework can simultaneously evaluate task-relevant and task-irrelevant variabilities when the averaged kinematics or task parameters change while considering both the motion sequence and task function.

Motor variability has another striking feature: the variability is embedded in a low-dimensional space that is referred to as synergy [6, 19-22]. It is suggested that to overcome a larger DoF, our motor system controls not all the DoFs but only those in low-dimensional space. Synergy has been (mainly) discussed for kinematics data [20, 22-24] and EMG data [19, 21]. In both kinematics and EMG, low-dimensional space that can capture a high portion of motor variability has been found. Several methods have been developed to extract the synergy, such as principal component analysis (PCA) [20, 22], nonnegative matrix factorization [21, 25] or spatial-temporal decomposition of EMG data [19].

The motor variability thus has at least two characteristics: compensation of the task-relevant variability and low-dimensional structure. Most of the techniques, however, deal with only one aspect. It is difficult to detect the low-dimensional structure of motor variability by the methods to evaluate task-relevant and task-irrelevant variabilities. Similarly, it is difficult to evaluate task-relevant and task-irrelevant variabilities by the techniques to extract low-dimensional structure of motor variability. Thus, the 1st principal component, the dimension that can explain the most significant portion of variability among all dimensions, is not always the most relevant or irrelevant to task performance. Although UCM has been used to extract synergy [8], the primary advantage of UCM is not the extraction of the low-dimensional structure but the evaluation of task-relevant and task-irrelevant variabilities. Although a few studies have focused on linear discrimination analysis (LDA) to discuss task-relevant low-dimensional space [26, 27], LDA enables only discrimination, e.g., success or failure of the movement [28], in contrast to TNC and GEM, which can discuss motor performance based on continuous performance value. In summary, few methods simultaneously quantify two striking features of movement variability.

Here, we propose a flexible and straightforward machine learning technique that can evaluate movement variability by unifying the advantages of previous techniques: our framework can evaluate not only task-relevant and task-irrelevant variabilities even when averaged kinematics or task parameters change (e.g., before, during, and after motor learning) while considering motion sequence and task function but also how each synergy is relevant to task performance by extending PCA. The current study relied on a ridge regression [29], a linear regression technique that is robust in the presence of measurement noise, has a definite relation to PCA, and can evaluate how the motion of each body part at each time is relevant to task performance in a data-driven manner [30]. Our technique can thus enable the identification of task functions in a data-driven manner without any explicit function such as the parabola or the forward kinematics. First, we formalize the dissociation of motion sequence data into task-relevant components and task-irrelevant components by extending a ridge regression. Second, we construct a novel experimental paradigm to discuss the relation of a motion sequence to task performance based on goal-directed and whole-body movements. We further discuss motor adaptation in the current experimental setting. Third, we validate the decomposition of motion sequence data into task-relevant and task-irrelevant components based on our experimental data. Fourth, we clarify the relation between ridge regression and PCA, a popular method to extract the low-dimensional space of motor variability. In particular, we analytically reveal how each principal component is relevant to performance in the ridge regression. We also validate the analytical calculations based on our experimental data. Finally, we apply our method to motion sequence data in whole-body and goal-directed movements before and after motor adaptation. Because our method enables to discuss the modulation of movement variability before and after motor adaptation, we discuss the dependence of the modulation on the perturbation schedule.

## Results

Downloading our program code on our website is possible.

### Linear regression

The current study relied on linear regression to determine the relationship between motion data ***X*** ∈ R^*T×D*^ (the current study focused on the temporal sequence of joint angles and angular velocities) and performance data ***d*** ∈ R^*T*×1^ following ***h*** = ***Xw***, where *T* and *D* denoted the number of trials and the number variables in the motion data, ***h*** ∈ R^*T*×1^ is the predicted performance, and ***w*** ∈ R^*D×*1^ is the best linear coefficients to predict the performance [30]. ***X***_*t*_, the *t*th row of ***X*** or the motion data at the *t*th trial, consists of vectorized motion data (e.g., after measuring joint angles of knee ***q***_k*,t*_ ∈ R^1*×F*^ and hip ***q***_h*,t*_ ∈ R^1*×F*^ for *F* time frames at the *t*th trial, ***X***_*t*_ = (***q***_k*,t*_, ***q***_h*,t*_)). We relied on a ridge regression to predict performance; the method enabled to predict the performance with higher accuracy than the conventional method as mentioned below. Ridge regression is a linear regression technique that is robust against observation noise, is applicable to data with multicollinearity (see *Materials and Methods* for details). Although we relied on a ridge regression to estimate ***w***, the following decomposition of input data into the task-relevant and task-irrelevant components could be applied to any linear regression technique.

### Decomposition into task-relevant and task-irrelevant components

After estimating the best linear coefficients ***w*** based on measured performance data ***y*** ∈ R^*T*×1^ and motion data ***X***, the estimated ***w*** enabled not only the prediction of performance but also the decomposition of motion data into a task-relevant component ***X***_rel_ and task-irrelevant component ***X***_irr_. By minimizing the cost function

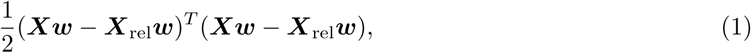

under the constraint ***X* ≠ *X***_rel_ to avoid the self-evident answer, ***X***_rel_ can be written as

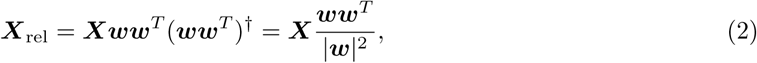

where (***ww***^*T*^)^*†*^ is a pseudo-inverse of ***ww***^*T*^ and 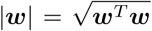. The equality 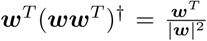 holds when ***w*** ∈ R^*D×*1^. Under the decomposition ***X*** = ***X***_rel_ + ***X***_irr_, ***X***_irr_ can be written as

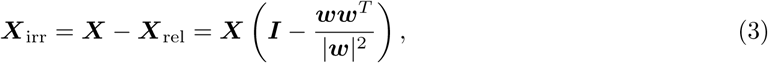

where ***I*** ∈ R^*D×D*^ is an identity matrix. Under the appropriate normalization (i.e., mean and standard deviation of each component of ***X*** and ***y*** were set to be 0 and 1, respectively, see *Materials and Methods* for details), ***X***_rel_***w*** = ***Xw*** = ***h*** and ***X***_irr_***w*** = 0, indicating that ***X***_rel_ and ***X***_irr_ denoted task-relevant and task-irrelevant components under the framework of linear regression. An important point of this framework is that it does not require any explicit function (e.g., forward kinematics such as in UCM or task function such as in GEM and TNC) but require only data ***X*** and ***y***.

Figs. 1A and 1B demonstrate typical examples of the decomposition when ***X*** includes only 2 elements and constrains the task by setting *h* = *X*_1_ − *X*_2_ (i.e., *w*_1_ = −1 and *w*_2_ = − 1) to some certain values (e.g., *y* = 2, 0, -2 in the simulated task 1, 2, 3, respectively). Because the constrained task was one dimensional and input data were two dimensional, an infinite patterns of ***X*** values resulted in an identical *h* value. In this case, 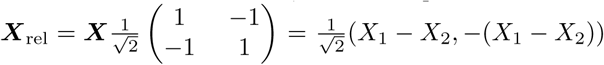 and ***X***_irr_ = ***X*** − ***X***_rel_. The simulated data points on the dotted line in Fig. 1B indicated ***X***_rel_. On the dotted line, the data points can be clearly separated into three parts corresponding to the simulated tasks 1, 2, and 3. In contrast ***X***_irr_ plotted on the solid line is not separated based on task.

**Figure 1.**
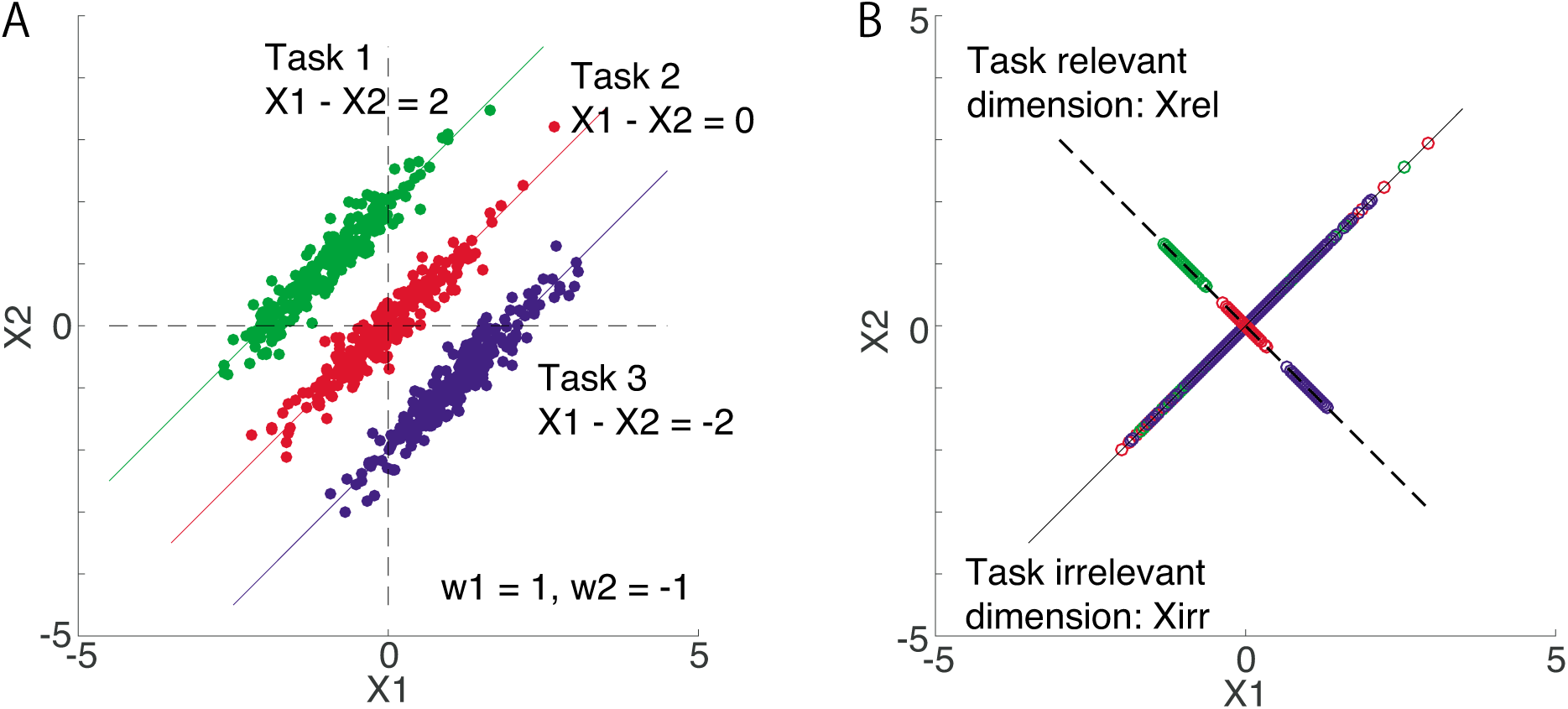
The concept of our method. **A**: An example of decomposing input data ***X*** into task-relevant ***X***_rel_ and task-irrelevant components ***X***_irr_. In this case, we assumed that the task 1 required *X*_1_ − *X*_2_ to be 2 (green line), task 2 required *X*_1_ − *X*_2_ to be 0 (red line), and task 3 required *X*_1_ − *X*_2_ to be -2 (blue line). Green, red, and blue dots indicate the typical input data for tasks 1, 2, and 3, respectively. In the ridge regression, these tasks can be achieved with *w*_1_ = 1 and *w*_2_ = −1, i.e., *h* = *w*_1_*X*_1_ + *w*_2_*X*_2_ = *X*_1_ − *X*_2_ should be determined differently in each task. **B**: The input data were decomposed into a task-relevant (black dotted line) component ***X***_rel_ = ***Xww***^*T*^ */|****w***|^2^ and a task-irrelevant component ***X***_irr_ = ***X*** − ***X***_rel_ (solid black line). ***X***_rel_ was separated depending on the task, and ***X***_irr_ was not separated, which indicates that the decomposition enables the discussion of the task-relevant and task-irrelevant components.

### Goal-directed whole-body movements and motor adaptation

The current study focuses on goal-directed and whole-body movements in which subjects manage to achieve the desired movement by controlling a large number of DoFs. We focus on a simplified version of whole-body movements: a vertical jump while crossing arms in front of the trunk (Fig. 2A). This goal-directed whole-body movement enabled us to focus on lower limb and trunk motions and assess task-relevant variability, task-irrelevant variability, and the low-dimensional spaces in which a high portion of the motor variability was embedded. We proposed a machine learning technique to simultaneously evaluate these features of variability while considering motion sequence and the relevance of each motion to jumping height.

**Figure 2.**
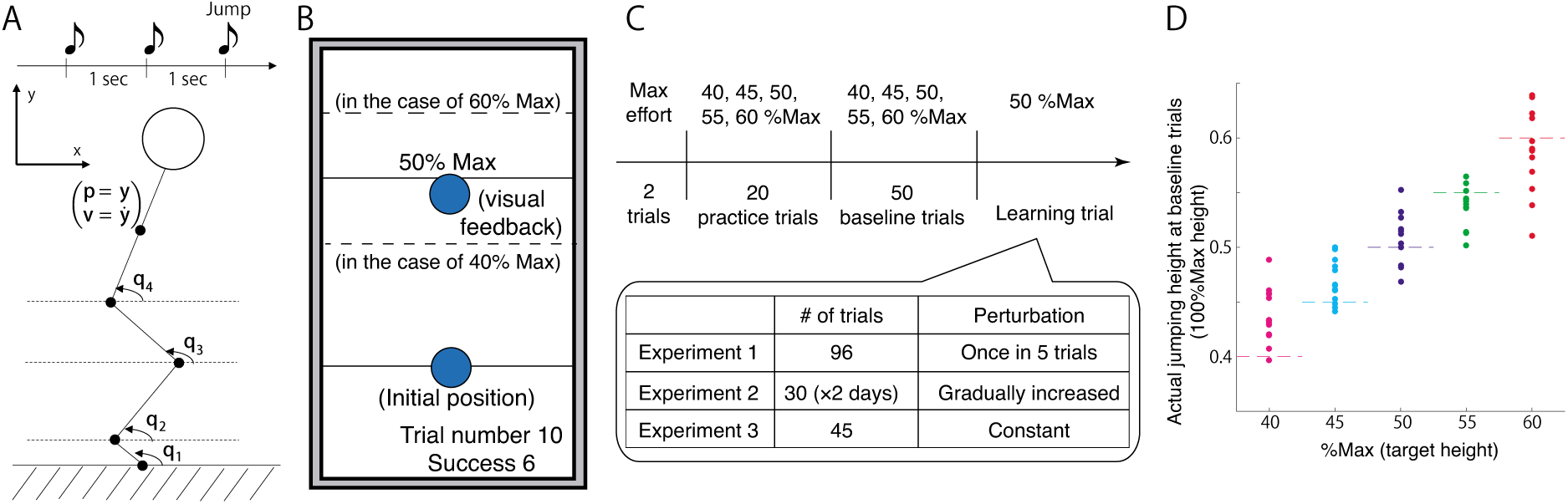
Summary of our experimental settings. **A**: Participants were instructed to perform a vertical jump at the timing of the third beep. Three beeps sounded at one-second intervals. We measured and analyzed joint angles at toe, ankle, knee, and hip in the sagittal plane. The jumping height was measured based on the position of the marker attached to the back in the y-axis. **B**: Task instruction and feedback information in each trial. A computer monitor was located in front of the participants (1.5 meters ahead, 1.7 meters above the floor). One second before the first beep, target height (indicated by black bar and texts [e.g., 50% max]), baseline height (indicated by black bar), and initial position (shown by blue cursor located on the baseline height) were displayed. When the target height was 60%, the black bar and text were presented at the position of the higher black dotted bar. When the target height was 40%, the black bar and text were displayed at the location of the lower black dotted bar. These black dotted bars were used only for the explanation and were not visible throughout the experiments. In the practice trials, the blue cursor was displayed during trials to continuously indicate the position of the marker attached to the back in the y-axis. These trials enabled the participants to become accustomed to the experimental setting. In baseline and learning trials, the blue cursor was displayed at the beginning and end of each trial. At the beginning of each trial, the blue cursor was presented at the baseline height. At the end of each trial, the cursor was displayed depending on the actual jumping height. When the jumping height was close to the target height, the participants heard a coin-getting sound. During the experiments, the subjects were provided with the current trial number and the number of successful trials. **C**: The sequence of the experiments. Participants performed a vertical jump with maximum effort for two trials. These jumping heights were used to determine the target height. Participants experienced 20 practice trials, 50 baseline trials, and a number of learning trials specific to each experiment. **D**: Averaged jumping height of each participant in the baseline trials in experiment 1. The jumping height depended on the target height (one-way repeated measure ANOVA, p = 6.114 × 10^−24^), indicating that the participants could perform the goal-directed movement.

Subjects stood in a fixed position and were instructed to look at a computer monitor located in front of them and perform a sub-maximum vertical jump according to a target height (40, 45, 50, 55, or 60% of the maximum jumping height of each subject, Fig. 2B, see *Materials and Methods* section for details). Three beeps sounded, and the subjects needed to perform the jump at the timing of the third beep. The interval between each beep was one second. At the beginning of each trial (i.e., one second before the first beep), the target height was indicated by a black bar displayed on a computer monitor. At the end of the *t*th trial, the actual jumping height *k*_*t*_ (the y-position of the marker attached to the back) was displayed as a blue cursor on the monitor, where *t* = 1*,…, T* and *T* was the number of trials to be analyzed. By manipulating the displayed jumping height (we called this manipulation as a perturbation *p*_*t*_), it was possible to induce sensory prediction error between the predicted and actual jumping heights. This perturbation paradigm was similar to a protocol of motor gain adaptation as reported mainly in saccade and arm-reaching movements [31, 32]. We expected subjects to modify their motion sequences to minimize the sensory prediction error.

First, we determined whether the subjects could perform goal-directed whole-body movements in our experimental setting. In 50 baseline trials in experiment 1 (Fig. 2C), the target height pseudorandomly changed in each trial. There was a significant main effect of target height in jumping height (Fig. 2D, one-way repeated measure ANOVA, p = 6.114 × 10^−24^). The subjects could thus perform goal-directed vertical jump depending on target height.

Second, we determined whether the subjects showed motor adaptation in the experimental setting. In 96 learning trials in experiment 1 (Fig. 2C), the subjects experienced perturbations once in every five trials; the perturbation was pseudorandomly set to *p*_*t*_ = 0.05 or *p*_*t*_ = − 0.05 in every five trials and *p*_*t*_ = 0 in other trials (Figs. 3A and B). We observed the modification of jumping height after each perturbation (Fig. 3C, paired t-test, *p* = 0.0026 for motor adaptation when *p*_*t*_ = 0.05; and *p* = 0.0014 for motor adaptation when *p*_*t*_ = − 0.05). Additionally, we confirmed whether fatigue influenced the adaptation by comparing the magnitudes of the modification in the former part of the learning trials to those in the latter part. There was no significant difference in the magnitudes of the modification between the former and the latter parts of the learning trials (paired t-test, *p* = 0.4382). Motor adaptation could thus be observed in the goal-directed vertical jump without significant effect of fatigue.

**Figure 3.**
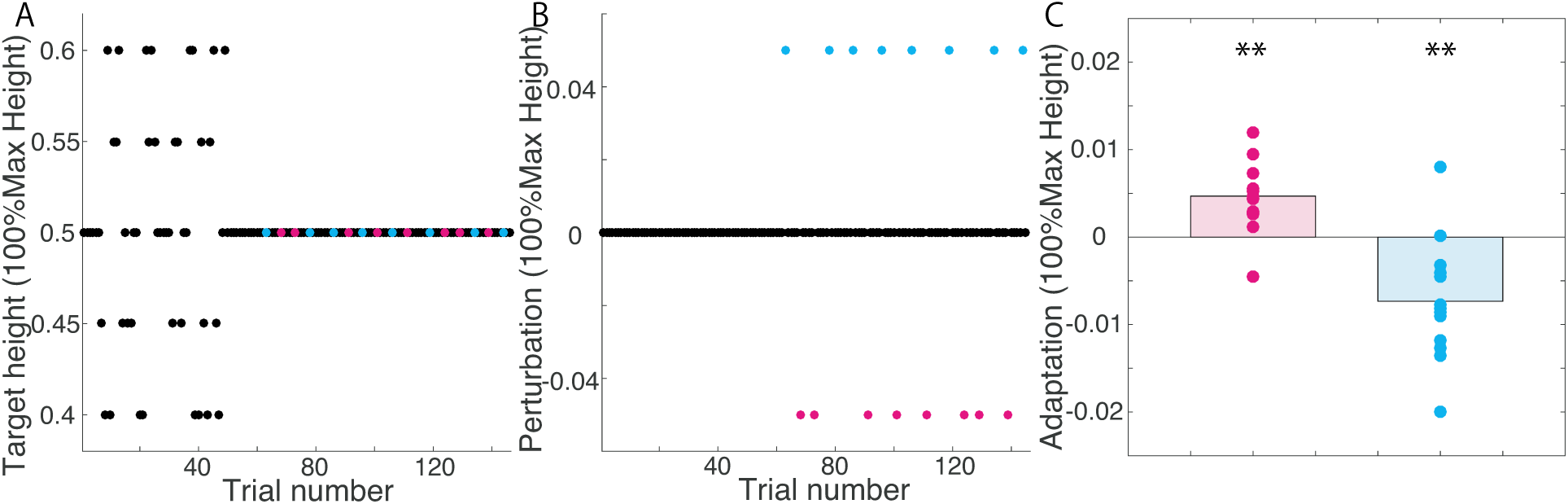
Diagram and results of experiment 1. **A**: Target height in baseline and learning trials. Cyan and magenta circles indicated the trials with perturbations. **B**: Perturbation sequence. The cyan circle indicates the trials with p=0.05, and the magenta circle indicates those with *p* = − 0.05. The perturbations were pseudorandomly imposed once in five trials. **C**: Adaptation effect. The vertical line indicates the modification of the jumping height after the perturbation was imposed. Magenta dots indicate the averaged difference in each subject corresponding to the perturbation *p* = − 0.05, and cyan dots indicate the averaged difference in each subject corresponding to the perturbation *p* = 0.05.

### Validation of ridge regression and decomposition into task-relevant and task-irrelevant components

The current study focuses on the evaluation of motor variability, especially task-relevant variability, task-irrelevant variability, and the relevance of low-dimensional structures to task performance, by extending ridge regression (the details of ridge regression were provided in *Materials and Methods*). We needed to validate the ridge regression in the current experimental setting before evaluating variability. Notably, we have already validated the efficiency of ridge regression to predict performance not only in jumping movements but also in throwing movements [30].

Ridge regression requires selecting input data because a careful selection of input data is indispensable to discussing the linear relation between input and output data. Prediction power is a sophisticated measure for selecting input data while avoiding overfitting [28]. The current study focused on prediction error between actual and predicted jumping height using 10-fold cross-validation. We compared the following three types of input data (see *Materials and Methods* for details). The first candidate is joint angles {*q*_*i*_} and angular velocities: 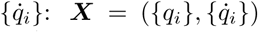, where {*a*_*i*_} = (*a*_1_*, a*_2_*, a*_3_*, a*_4_) and *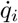* denotes the derivative of *q*_*i*_ concerning time (the definitions of each *q*_*i*_ are given in Fig. 2A). The second candidate is the functions of *q*_*i*_ and 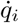, which describe the position and velocity of back joint in the y-axis and are relevant to jumping height: 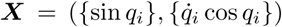. The third candidate is the functions describing the jumping height based on parabolic approximation: 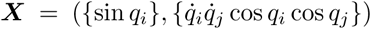, where 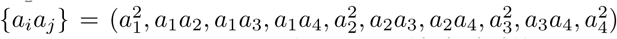. By comparing these three candidates, we found that the first candidate (i.e., 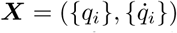) showed the lowest prediction error (Fig. 4A). In particular, the first candidate, with four-time frames before release, yielded the lowest prediction error. If the prediction error equals 1, the method cannot predict the output data. In contrast, if the prediction error equals 0, the method can predict the output data with 100% accuracy. As shown in Fig. 4A, the first candidate with four-time frames resulted in a prediction error of 0.174, indicating that the ridge regression enables prediction of jumping height with an 82.6±2.28% (± mean standard error of the mean [s.e.m.], N=13) accuracy in the current setting. In the following, we refer to the first candidate with four-time frames as the motion sequence. In our experimental setting with goal-directed vertical jump, ***X*** included 32 elements in each trials (4(dim)×4(time frames) for {*q*_*i*_}, and 4(dim)×4(time frames) for 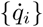. Because the parabolic approximation of jumping height (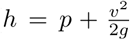, the detailed definition is given in the *Introduction*) enables prediction of the jumping height with a 76±2.96% (mean±s.e.m., N=13) accuracy (purple line in Fig. 4A), the ridge regression enables prediction of jumping height with higher accuracy than the approximation. The reasons the ridge regression shows higher prediction power are its robustness against observation noise and consideration of the motion sequence rather than the representative motion data at a single time frame (i.e., the position and velocity of the hip joint only at the time of release). The ridge regression thus enables the discussion of the linear relation between the motion sequence and jumping height with appropriate precision.

**Figure 4.**
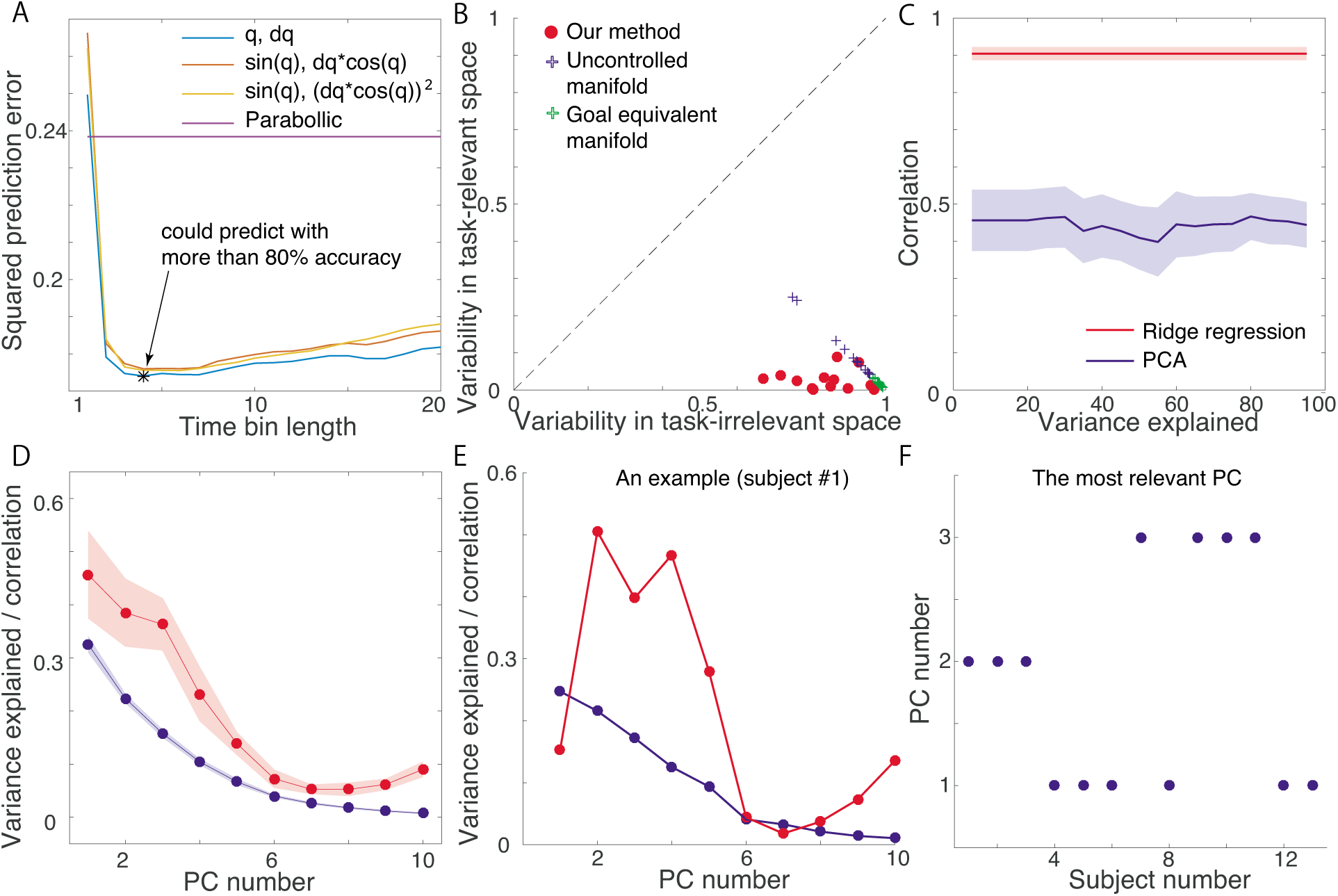
Validation of our method and comparison to previous methods. **A**: Predictive power of the ridge regression using three kinds of input data. Horizontal and vertical axes indicate the time bin length used for the ridge regression and squared prediction error, respectively. If the ridge regression could not make a prediction, the prediction error equaled 1. If the ridge regression could predict the output data perfectly, the prediction error equaled 0. These results indicate that ridge regression enables the prediction of output data with an 82.6 *±* 2.28% (mean *±* s.e.m., N=13) accuracy. **B**: Evaluation of task-relevant and task-irrelevant variabilities. Red dots indicate those variabilities evaluated by our method in each subject (N = 13). Blue and green crosses indicate the variabilities evaluated by the UCM and GEM, respectively. Our method uses a different normalization method from those of the UCM and GEM. **C**: Correlation between predicted and actual jumping height. The red line and the shaded area indicate the mean and standard error of the mean (s.e.m.) of the correlation in ridge regression (N = 13), respectively. The blue line and shaded area indicate the mean and s.e.m. of the correlation in PCA (N = 13), respectively. Horizontal and vertical axes indicate the explained variance or the corresponding number of principal components and the correlation, respectively. **D**: Explained variance and the correlation between the predicted and actual jumping height of each principal component. The red line and the shaded area indicate the mean and s.e.m. of the correlation (N = 13), respectively. The blue line and the shaded area indicate mean and s.e.m. of the variance explained (N = 13), respectively. **E**: Variance explained and the correlation between the predicted and actual jumping height of each principal component in a typical subject. The red and blue lines indicate the correlation and the variance explained, respectively. **F**: The PC number that is most relevant to predicting jumping height. Horizontal and vertical axes indicate the subject number and the most relevant PC number, respectively.

### Variability in task-relevant and task-irrelevant space

We calculated the task-relevant and task-irrelevant variabilities in a goal-directed vertical jump based on both the ridge regression and the decomposition of input data into task-relevant and task-irrelevant dimensions. The current study calculated the variability (variance) of each element of ***X***_rel_ and ***X***_irr_ in focused trials (see *Materials and Methods* for details). The representative values of the variability, task-relevant variability Var_rel_ and task-irrelevant variability Var_irr_, were calculated by averaging the variability across all the elements.

We found that the task-relevant variability was smaller than the task-irrelevant variability in all participants (N = 13, red dots in Fig. 4B). Because previous methods, such as the UCM (blue crosses) and GEM (green crosses), found similar task-relevant and task-irrelevant variabilities, our method enabled the extraction of lower task-relevant variability and higher task-irrelevant variability. Because the normalization procedures of our method and previous methods differ (see *Materials and Methods*), there was a slight difference in the calculated variabilities. Our methods can quantify task-relevant and task-irrelevant variabilities by considering the motion sequence and the relevance of the sequence to the task. Our method does not require any explicit task function, such as the parabolic approximation of jumping height, but it determines the relevance of the motion sequence to the task in a data-driven manner. Further, our method is robust against observation noise due to the properties of ridge regression.

### Relevance of each principal component to task performance

Movement variability shows not only less task-relevant variability than task-irrelevant variability but also a low-dimensional structure. The current study compares our method to PCA, a conventional method to extract low-dimensional structure. Because the low-dimensional structure is considered to represent some features of motor control, it can be expected to be correlated to task performance. We decomposed the motion sequence ***X*** into principal components (PCs, i.e., eigenvectors) and calculated the correlation of each PC to jumping height (see *Materials and Methods* for detail). In our setting, there was no clear relation between the number of PCs for the decomposition and the correlation between the decomposed motion data and performance data (Fig. 4C). If averaged across all participants, the 1st PC could explain approximately 30% of the movement variability (blue line in Fig. 4D). Corresponding to the explained movement variability, the 1st PC showed the highest correlation to jumping height (red line in Fig. 4D) if averaged across all participants. In a typical subject, however, the 2nd rather than the 1st PC showed the highest correlation to jumping height (red line in Fig. 4E). This typical subject was not an exception; Fig. 4F shows the PC number with the highest correlation to jumping height. In 6 out of 13 subjects, the 1st PC showed the highest correlation to performance. In 3 out of 13 subjects, the 2nd PC showed the highest correlation, and the 3rd PC showed the highest in 4 out of 13 subjects. These results indicate that the explained movement variability did not correspond to the relevance to task performance.

Ridge regression enabled the prediction of jumping height with higher accuracy than PCA (red line in Fig. 4C) because the ridge regression weights each PC based on both the explained movement variability and the task relevance. In PCA (or equivalently singular value decomposition (SVD)), the motion sequence at the *t*th trial is decomposed as 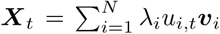, where *N* is the number of PCs, *λ*_*i*_ is the *i*th eigenvalue corresponding to the *i*th PC ***v***_*i*_, and *u*_*i,t*_ indicated how the *i*th PC appeared at the trial. The correlation of the *i*th PC to task performance was thus calculated based on *u*_*i,t*_, and it did not reflect the relevance of the *i*th PC to task performance. In contrast, the ridge regression enables the prediction of task performance as 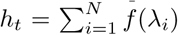 Corr(*u*_*i,t*_*, y*_*t*_)*u*_*i,t*_, where *f* (*λ*_*i*_) is a sigmoidal function of *λ*_*i*_ and Corr(*u*_*i,t*_*, y*_*t*_) is the correlation between the contribution of the *i*th PC at the *t*th trial *u*_*i,t*_ and observed jumping height *y*_*t*_ (see *Materials and Methods* for details). The ridge regression thus enables the prediction of performance by weighting each PC based on both explained movement variability and task relevance. In other words, our method enables the consideration of the low-dimensional structure of movement variability by weighting each PC suitable for predicting task performance.

### Influence of motor adaptation on variability in the task-relevant and task-irrelevant dimension

An advantage of our method is its linearity, which enables the simultaneous comparison of the task-relevant and task-irrelevant variabilities among the conditions where mean kinematics or task parameters change (e.g., before, during, and after motor learning). It was previously unclear how task-relevant and task-irrelevant variabilities are modulated by motor adaptation. The modulation of these variabilities has been investigated for arm-reaching movements and motor adaptation to a constant perturbation [5, 33]. Although there are some differences between adaptation to a constant perturbation and that to a gradually imposed perturbation, e.g., retention rate or awareness [34], the means by which those variabilities are modulated in the two types of adaptations have not been investigated. Further, it was previously unclear whether such modulation of variability could be observed in whole-body movements. Our method without linear approximation enabled the discussion of how task-relevant and task-irrelevant variabilities are modulated before and after motor adaptation in whole-body movements. We thus applied our method to motor adaptation in response to constant and gradually imposed perturbations.

In experiment 2 (two days for each subject), subjects experienced gradually increased or decreased perturbations. Each subject underwent ten learning trials without any perturbation. The perturbation was gradually imposed for ten trials and was set to 0.05 or -0.05 for ten trials (Figs. 5A, B). The gradually imposed perturbation required not abrupt but gradual compensation (i.e., subjects were required to modify their motions slightly in each trial). In a total of 30 trials, the target height was set to 50% of the subject’s maximum jumping height. Subjects who experienced a *p*_*t*_ *>* 0 on the first day experienced a *p*_*t*_ < 0 on the 2nd day and vice versa. The order of perturbation was counterbalanced across subjects. The subjects could adapt to the gradually increased or decreased perturbations (Fig. 5C).

**Figure 5.**
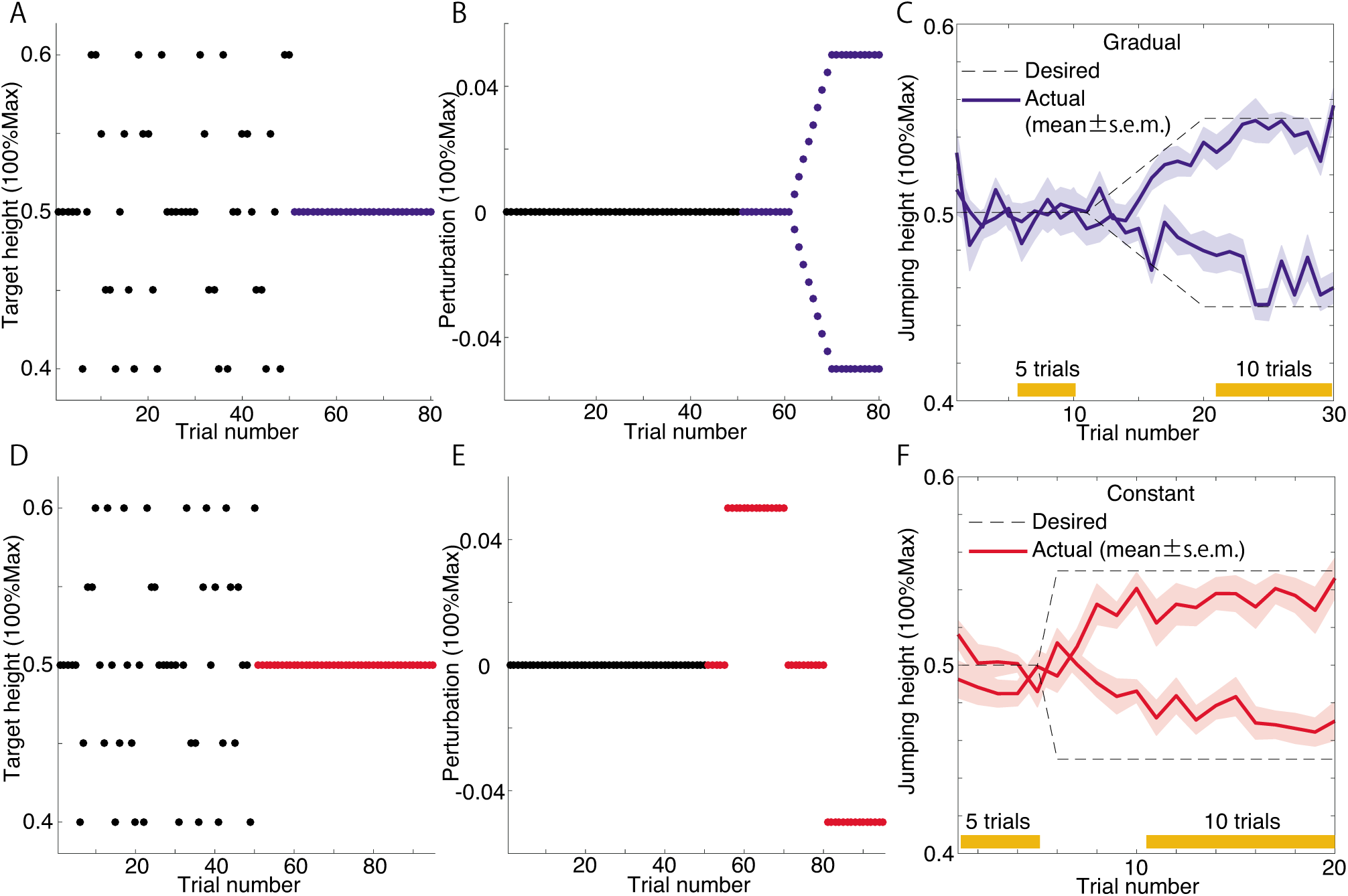
Diagram and results of experiments 2 and 3. **A,D**: Target height in baseline and learning trials. Blue and red circles indicate the trials with perturbations. **B,E**: Perturbation sequence. Subjects joined experiment 2 for two days and experienced two different perturbations (either *p >* 0 or *p <* 0). The order of perturbation was counterbalanced across subjects. In experiment 3, subjects experienced both positive and negative perturbations within one day. Although panel E showed the case when a negative perturbation followed a positive one, the order of the perturbation was counterbalanced across subjects. **C,F**: Learning curves. The thin solid lines indicate the learning curves of each subject. The bold solid line indicates the learning curve averaged across all subjects.

In experiment 3, the subjects experienced constant perturbations. Each subject underwent five learning trials without any perturbation. The perturbation was set to 0.05 or -0.05 for 15 trials, 0 for ten trials for washout and -0.05 or 0.05 for 15 trials (Figs. 5D, E). In contrast to experiment 2 where the perturbation was gradually imposed, subjects were required to modify their motions abruptly in experiment 3. Subjects who experienced a *p*_*t*_ = 0.05 on the 6th-20th trials experienced a *p*_*t*_ = − 0.05 on the 31st-45th trials and vice versa. The order of perturbation was counterbalanced across subjects. In a total of 45 trials, the target height was set to 50% of the subject’s maximum jumping height. In both experiments 2 and 3, the subjects adapted to the perturbations (Fig. 5F).

We calculated the task-relevant and task-irrelevant variabilities before and after adaptation in experiments 2 and 3 (Figs. 6A and B). For task-relevant variability, there was no significant difference before and after the adaptation to gradually increasing or decreasing perturbations (blue dots in Fig. 6A, N = 13, Wilcoxon signed rank test, p = 0.1909). In contrast, in adapting to a constant perturbation, there was a significant difference in task-relevant variability before and after adaptation (red dots in Fig. 6A, N = 13, Wilcoxon signed rank test, p = 0.0034). For task-irrelevant variability, there was no significant difference before and after adaptation to gradually increasing or decreasing perturbations (blue dots in Fig. 6B, N = 13, Wilcoxon signed rank test, p = 0.1677) and a constant perturbation (red dots in Fig. 6B, N = 13, Wilcoxon signed rank test, p = 0.3396). These results could be interpreted based on a simulated and two-dimensional case similar to that shown in Fig. 1 (Figs. 6C, D). In adapting to perturbations, the subjects needed to modify their output (i.e., jumping height) by determining an appropriate input (i.e., motion sequence). In adapting to gradually increasing or decreasing perturbations, there was no modulation in both task-relevant and task-irrelevant variabilities (Fig. 6C). In adapting to a constant perturbation, the task-relevant variability increased, while the task-irrelevant variability was not be modulated (Fig. 6D). Notably, Figs. 6C and 6D were not real data but simulated examples to interpret our results. In summary, the modulation of task-relevant variability depends on the schedule of perturbation.

**Figure 6.**
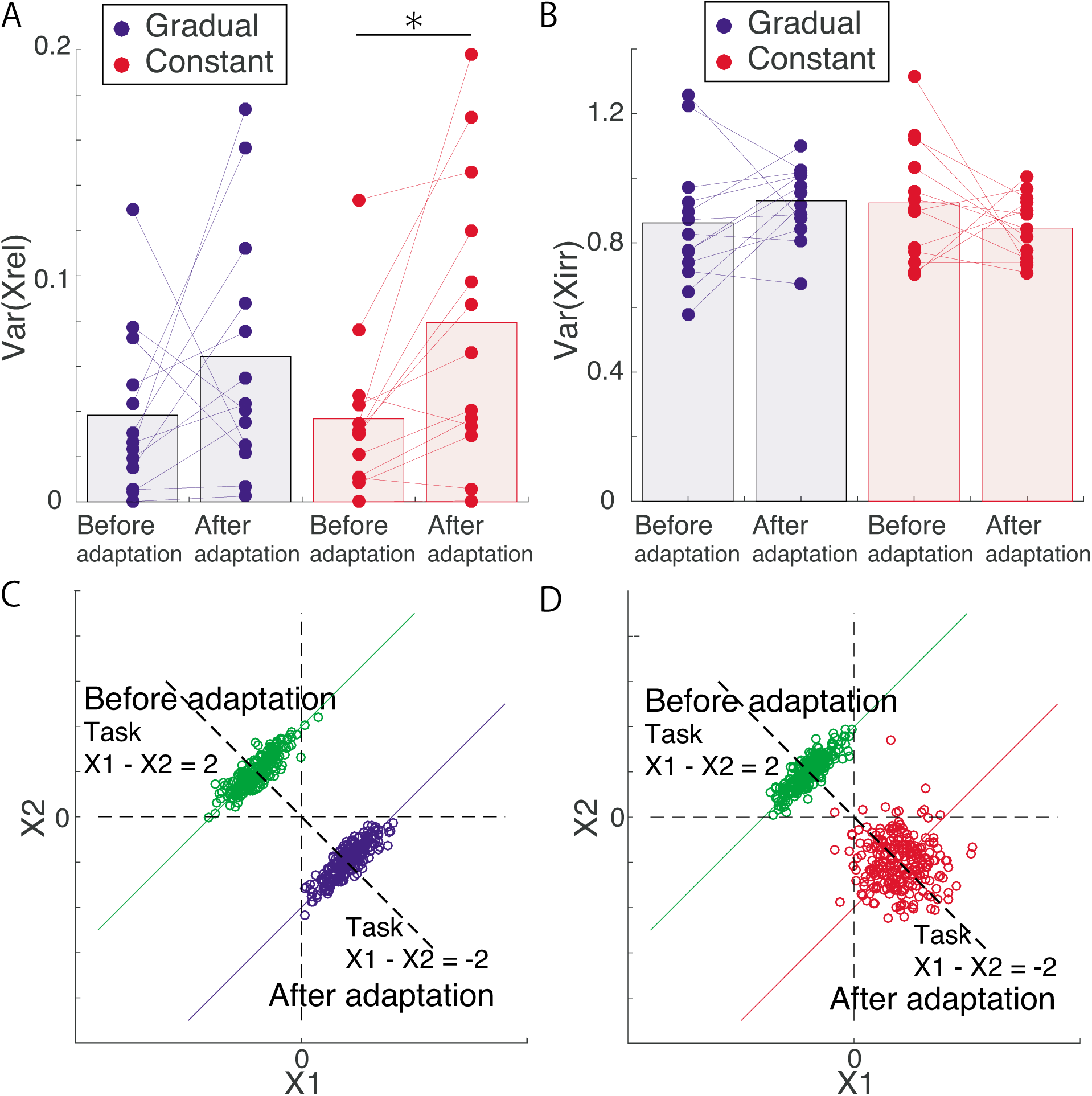
Application of our method to the results of experiments 2 and 3. **A**: Task-relevant variabilities of each subject (N = 13) before and after adaptation to perturbation in experiments 2 and 3. The blue and red dots indicate the variability of each subject in experiments 2 and 3, respectively. The blue and red lines show the modulation of the variability due to adaptation. The blue and red bars indicate the averaged variability across all subjects. There was a significant difference between the variabilities before the adaptation and those after the adaptation in experiment 3 (Wilcoxon signed rank test, p = 0.0034). **B**: Task-irrelevant variabilities of each subject (N = 13) before and after adaptation to perturbation in experiments 2 and 3. **C,D**: Interpretation of our results based on a simple example. We assume that the task before adaptation required *X*_1_ − *X*_2_ to be 2 and that the task after adaptation required *X*_1_− *X*_2_ to be -2. Panel C indicates an interpretation of our results in experiment 2. In experiment 2, there was no modulation in both task-relevant and task-irrelevant variabilities. Panel D suggests an explanation of our findings from experiment 3. In the experiment, task-relevant variabilities increased after adaptation, and task-irrelevant variabilities remained unchanged.

## Discussion

We proposed a flexible and straightforward machine learning technique that quantified task-relevant variability, task-irrelevant variability, and the relevance of each principal component to task performance in a noise-robust manner while considering motion sequence and how each motion sequence was relevant to task performance (Fig. 4). Our method can find the relevance of each motion sequence to performance (i.e., task function) in a data-driven manner; our method does not require any explicit task function, such as the parabolic approximation of jumping height. Further, our method does not require any linear approximation, which enables the simultaneous consideration of the variabilities when the kinematics or task parameters averaged across trials change (e.g., before, during, and after adaptation). By applying our method to the motion sequence before and after motor adaptation, we found that the perturbation schedules affected the modulation of movement variability in motor adaptation (Figs. 6A and 6B). These advantages enable the methods to be flexibly applied to a wide range of goal-directed movements.

Our method can be regarded as a generalized method including UCM and GEM. When we define 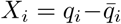 (*q*_*i*_ and 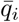 indicated the *i*th joint angle and the averaged joint angle across all the focused trials, respectively [i=1, …, 4 in our setting]), and 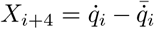 (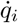 and 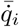 indicated the joint angular velocity and averaged joint angular velocity across all the focused trials, respectively), and the corresponding weight ***w*** as the Jacobian matrix of forward kinematics *p* = *p*(***q***) (the back position) and 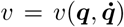 (the back velocity) around the averaged joint angles and angular velocities across all the focused trials 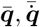, our framework corresponds to UCM. When we define 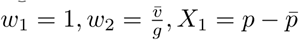, and 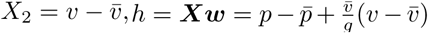, our framework corresponds to the GEM in the cases when the task function can be defined by the parabolic function, where *g* denotes the gravitational acceleration. Because the UCM and GEM can be regarded as a special case of our method, our method can be considered as a generalized version of those methods.

Another advantage of our method is the ability to select appropriate input based on predictive power (Fig. 4A). The predictive power also enables the selection of a proper coordinate to define the task performance. A previous study has demonstrated that the UCM and TNC frameworks are sensitive and insensitive to how to select the coordinate (e.g., either relative or absolute angle), respectively [35]. Our framework is likely sensitive to how to choose the coordinate; however, in contrast to the UCM framework, our method enables the selection of the appropriate coordinate for discussing the relationship between motion and performance based on predictive power. Although we considered one-dimensional performance in the current study (i.e., jumping height), two-dimensional performance requires the definition of an appropriate coordinate to discuss performance [30, 36]. Predictive power plays a vital role in selecting the proper coordinate not only in motion but also in the performance space [30]. How to select the length of time frames is another crucial problem (Fig. 4A). In our data, the motion data with four-time frames (approximately 33 ms) was chosen for the best predictive power. Although the motion data with four-time frames sound motion fragments rather than motion sequences, our method can be applied independently of the length of time frames. In our case, four-time frames were chosen to increase the predictive power.

Although we compared our method to UCM and GEM (Fig. 4B), we needed to compare it to the TNC method [13,37], the other method used to quantify motor variability from a different perspective. TNC enables the extraction of three types of information from motion data: T-cost, which quantifies how the mean motion data deviate from the optimal motion; N-cost, which quantifies how the motor variability deviates from the optimal variability; C-cost, which quantifies how the covariance among motion data deviates from the optimal covariance. Although the TNC does not consider task-relevant and task-irrelevant variabilities, it can quantify other interesting features (i.e., T-, N-, C-costs) embedded in motion data and variations of those costs during the learning process. Due to the computational cost, we could not apply the TCN method to our data. The TCN requires a grid search in the calculation of T-cost. Because the number of the grid was 200 and that of focused variables was eight in our case (four joint angles and angular velocities), it required 200^8^ calculations. Due to this too burdensome computational cost in calculating not only T-cost but also C-cost, we could not apply the TNC method to our case. When the number of focused variables is two, the TNC could be a promising method and work well [13,37].

A potential extension of our method is to yield a state-space model for motor adaptation during whole-body movements. The state-space model was previously proposed as a model of motor adaptation mainly in arm-reaching movements [38-46]. In the current study, the modification of jumping height at the *t*th trial, *h*_*t*_ − *h*_*t*−1_, was significantly correlated with the error, *e*_*t*−1_, caused by perturbation and motor noise (in experiment 1, the correlation between *h*_*t*_ − *h*_*t*−1_ and *e*_*t*−1_ averaged across all participants was 0.5118 and *p <* 0.01 for all the participants). The state-space model of jumping height can thus be written as *h*_*t*_ = *h*_*t*−1_ + *ηe*_*t*−1_, where *η*(*>* 0) is the learning rate. The jumping height *h*_*t*_ was predicted well by *h*_*t*_ *≃* ***X***_*t*_***w*** = ***X***_rel,t_***w***, which enabled us to approximately rewrite the state-space model as ***X***_rel,t_***w*** = ***X***_rel,t−1_***w*** + *ηe*_*t*−1_. The model indicated that the jumping height was modified via modifying the motion sequence in the dimension along ***w***. Because this is a possible future extension of our approach, we need to further investigate the above-mentioned frameworks.

An advantage of our method is the linearity (i.e., ***h*** = ***Xw***) in contrast to nonlinearity inherent in our body dynamics. A likely explanation for why linear regression works well is by analogy with the motor primitive framework, a successfully used framework in motor adaptation with goal-directed arm-reaching movements [38-46]. In this framework, a nonlinear motor command *u* is modeled as the linear weighted sum of nonlinear neural activities ***A***; *u* = Σ_*i*_ *W*_*i*_*A*_*i*_ and *W*_*i*_ are modified to minimize the movement error between the actual hand position and the desired movement position. When *A*_*i*_ is a nonlinear function of the desired movement and appropriately high-dimensional, nonlinear motor commands can be generated by appropriate linear combinations of nonlinear neural activities, which has been theoretically validated in the framework of a basis function network [47]. Motion data ***X*** can be a nonlinear function of movement performance because our body dynamics are nonlinear. Additionally, the motion data are appropriately high-dimensional (32 dimensions for 1-dimensional performance). Thus, an appropriate linear summation ***Xw*** could predict the actual movement performance, which resulted in an appropriately estimated ***w*** that represented the relevance of motion elements to performance.

We relied on a simple linear regression (i.e., ridge regression). It is possible to use a more complicated machine learning technique, such as a mixture model [28, 48-50], sparse regression technique [51], or non-linear regression technique [52]. We have shown that a nonlinear regression technique, such as Gaussian process regression, is not effective in predicting performance based on motion data [30], likely because the number of data is limited. Although sparse regression, nonlinear regression, or a mixture model can show better predictive performance if the number of the data is high enough in general, it is difficult to find certain relations between the principal components and estimated parameters via those methods. Ridge regression enables the determination of not only task-relevant and task-irrelevant variabilities but also the relevance of each PC to performance. Because some previous studies have discussed the relevance of each PC to performance [53,54], it could be a promising research topic to evaluate the functional roles of low-dimensional structured from a different viewpoint.

To our knowledge, only a few studies have investigated how variability is modulated through motor adaptation [5, 33]. A previous study clarified that the variability is modulated after motor adaptation by utilizing a constant force field [33]. To our knowledge, it was not clarified whether the perturbation schedule affected the modulation. The current study suggests that the perturbation schedule changes the modulation of variability (Figs. 6A and B). Because the variability can facilitate exploration [33], the current study also suggests that constant perturbations facilitate exploration in task-relevant space and that gradually applied perturbations do not affect the exploration. Recent studies have suggested that the variability plays essential roles in sports performance [55], injury prevention [56], and the development of children with developmental coordination disorder [57]. The current result provides a hint about how to assist those functions through the facilitation of the exploration using false feedback.

## Materials and Methods

### Participants

Thirteen healthy volunteers (aged 18-22 years, two females) participated in all of our experiments. On the first day, the participants underwent ten practice trials and 160 baseline trials with pseudorandomly changing targets (40%, 45%, 50%, 55%, or 60% of the maximum jump height) and became accustomed to the experimental setting. At the second, third, fourth, and fifth days (not consecutive), they joined experiments 1, 2, and 3. They joined experiment 2 for two days. All participants were informed of the experimental procedures and their confirmation with the Declaration of Helsinki, and all participants provided written informed consent before the initiation of the experiments. All procedures were approved by the ethics committee of the Tokyo University of Agriculture and Technology.

### Data acquisition and processing

Jumping motions were recorded at 120 Hz using six cameras (Optitrack Flex 13, NaturalPoint Inc., Corvallis, Oregon). Markers were attached to the back (TV10), right hip joint (Femur greater Trochanter), right knee (Femur Lateral Epicondyle and Femur Medial Epicondyle), right heel (Fibula Apex of Lateral Malleolus and Tibia Apex of Medial Malleolus), and right toe (Head of 2nd Metatarsus) of the participants. Marker position data were filtered using a 12th-order, 10 Hz zero-phase Butterworth filter using MATLAB 2016a. Joint angles between the right toe and heel (*q*_1_), right heel and shank (*q*_2_), right shank and thigh (*q*_3_), and right thigh and trunk (*q*_4_) were calculated in the sagittal plane (Fig. 2A). Because the current study focused on a vertical jump while crossing arms in front of the trunk, it was possible to focus only on lower limb and trunk motions. Throughout the current study, we focused on the four-link model of the lower limbs in the sagittal plane.

Release timing was detected based on the moment at which the vertical toe position exceeded 10% of the maximum height in each trial. The predictive power was calculated using various time-bin lengths including the release timing (Fig. 4A). When the time bin length was four, the fourth time frame corresponded to the release timing, the third time frame corresponded to one time frame before the release timing, and the other time frames followed accordingly.

### Experimental setup

At the beginning of each trial, the subjects were instructed to stand at a fixed position. In each trial, subjects listened to three beeps separated by one-second intervals; the first beep indicated the start of each trial, and the subjects were required to jump at the timing of the third beep.

We measured the position of the marker attached to the subject’s back using MATLAB at 30 Hz. In front of the subject (1.5 meters ahead, 1.7 meters above the floor), there was a monitor to display a blue cursor that indicated the height of the marker attached to subject’s back and a black bar that indicated target height (Fig. 2B). Those cursors and bars were displayed one second before the first beep sounded. The blue marker could move only in the vertical axis because the current study focused on the vertical height of the jumping motion. The marker position at time *s* in the y-axis (Fig. 2A), *k*_*s*_, was displayed on the monitor after being normalized for each subject as 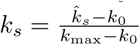, where 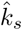 is the marker position without normalization, *k*_0_ indicates the initial marker position that was evaluated at an upright standing position in each trial, and *k*_max_ indicates the jumping height with maximum effort in each subject (Fig. 2C). For the two trials that required the subjects to jump with maximum effort, there was no cursor feedback. One second before the first beep, *k*_*s*_ = 0, the blue circle was displayed on the black baseline on the monitor. Additionally, the target height *d* was indicated by a black line. Before the baseline trials, the subjects underwent ten practice trials. In those trials, the marker position in each time frame was displayed on the monitor and *d* was pseudorandomly chosen from 0.40, 0.45, 0.50, 0.55, or 0.60 (each value was randomly chosen only once in every five trials). This method enabled the subjects to become acquainted with the experimental setting by confirming the motion trajectory of the marker attached to their back. In baseline trials, the marker position was displayed only at the start and end of each trial. One second before the first beep, the cursor was displayed on baseline position, and a black line was corresponding to *d* was displayed as pseudorandomly chosen from 0.40, 0.45, 0.50, 0.55, or 0.60. At the end of each trial, the cursor was displayed at the maximum value of *k*_*s*_ within each trial (i.e., max *k*_*s*_), which indicated jumping height (Fig. 2B). When the subjects achieved a jumping motion that was close to the target height (|*d*− max *k*_*s*_| < 0.02), they heard a coin-getting sound to indicate that the jumping motion was successful. After the baseline trials, the subjects underwent 96 learning trials in experiment 1, 30 trials in experiment 2 (the same set of practice and main trials was imposed for two days), and 45 trials in experiment 3.

We utilized a perturbation paradigm to investigate how subjects modify their jumping motion via experiencing sensory prediction errors. For trials with perturbation *p*, the position of the cursor was displayed at max *k*_*s*_ + *p*. The subjects needed to modify their jumping motion to achiever a lower (when *p >* 0) or higher jumping height (when *p <* 0). When the displayed jumping height was close to the target height (|*d*− (max *k*_*s*_ + *p*)| < 0.02), the subjects heard a coin-getting sound to indicate that the jumping motion was successful.

### Task-relevant and task-irrelevant variabilities

Under the condition ***X*** = ***X***_rel_ +***X***_irr_ (see *Results* for details), the variance of the *i*th component of ***X***, *X*_*i*_, can be calculated as

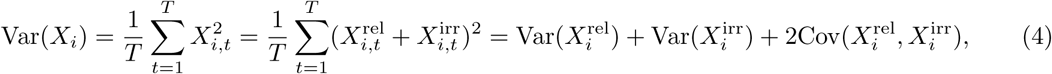

where *X*_*i,t*_ is *X*_*i*_ at the *t*th trial, 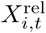 is the *i*th component of ***X***^rel^ at the *t*th trial, 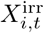 is the *i*th component of ***X***^irr^ at the *t*th trial, and 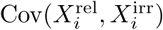 is the covariance between 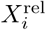 and 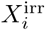 Notably, in the current experimental setting, 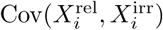 in the analyzed trials was close to 0. We thus considered only 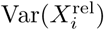 and 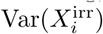

### Ridge regression

The ridge regression enabled us to determine the best one-dimensional linear space ***w*** ∈ R^*D×*1^ in the input data ***X*** ∈ R^*T×D*^ to predict the output data ***y*** ∈ R^*T*×1^ by minimizing the cost function:

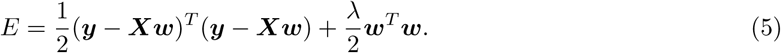

The first term on the right-hand side indicates the fitting error, the second term indicates the regularization of ***w***, and *λ* is a regularization parameter. The current study determined *λ* to minimize the prediction error based on a 10-fold cross validation, which enabled us to avoid overfitting [28]. Overfitting, which can appear without any regularization, leads to the selection of a model that is more complicated than the true one. Minimization of the cost function concerning ***w*** leads to the optimal value for ***w***:

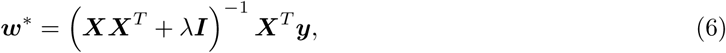

where ***I*** was an identity matrix. When ***XX***^*T*^ has multicollinearity, it is difficult to calculate the inverse of ***XX***^*T*^ because of the rank deficit. The identity matrix with a regularization parameter *λ* enables to calculate the inverse of ***XX***^*T*^ + *λ****I*** and predict output data with a certain accuracy.

The ridge regression showed high prediction power under the existence of measurement noise in ***X***. Under the existence of measurement Gaussian noise ***ξ*** with a mean of 0, the standard deviation is *σ*_*o*_, covariance is 0, and the cost function averaged across all the possible noise can be written as

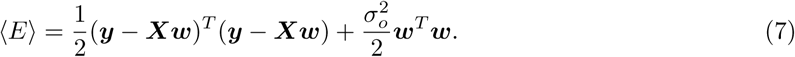

The equivalence between equations (5) and (7) indicates that the ridge regression enabled the selection of the best ***w*** to predict ***y*** under the existence of measurement noise while avoiding overfitting. The equivalence also suggests that the regularization parameter *λ* corresponds to the variance of the observation noise 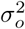.

The ridge regression enabled the estimation of an appropriate ***w*** based on the normalized ***y*** and ***X***, i.e., the mean and standard deviation of ***y*** and ***X*** should be normalized to be 0 and 1, respectively; 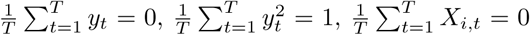, and 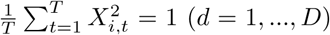 All the results in the current study depended on the normalized data. Without normalization, *w*_*i*_ is estimated to be large when *X*_*i,t*_ shows small fluctuations and vice versa, although regularization with parameter *λ* was imposed equally to all the *w*_*i*_; therefore, normalization, especially in ***X***, is indispensable for estimating appropriate ***w***. Notably, the normalization did not affect interpretation at all because it was possible to restore the original unnormalized data by adding the original mean 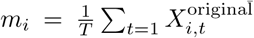 and multiplying the original standard deviation 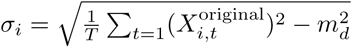 To satisfy 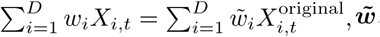, where ***w*** corresponds to unnormalized data, should be divided by 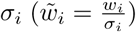 and 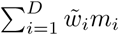 should be subtracted. In total, the normalization is indispensable for estimating an appropriate ***w***; however, it did not affect the results at all.

### Parabolic representation of jumping height, three candidates of input data, the UCM and GEM

The vertical position of the marker attached to the subject’s back determined the jumping height in the current study. We expected that the jumping height could be predicted well based on the back position *p* and velocity *v* of the marker at the release timing as follows

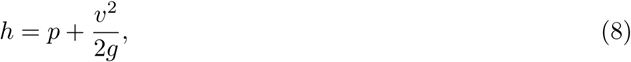

where *g ≃* 9.8(*m/s*^2^). In the joint angle representation, *p* and *v* were written as follows:

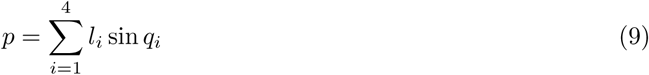

and

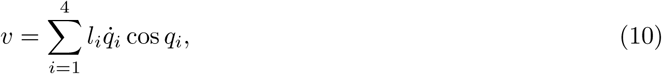

where *l*_*i*_ indicated the length of the *i*th limb (i.e., *l*_1_ indicated the length between right toe and heel, *l*_2_ indicated the length between right heel and knee, *l*_3_ indicated the length between right knee and hip, and *l*_4_ indicated the length between hip and back). In the UCM (blue crosses in Fig. 4B), we calculated the task-relevant and task-irrelevant variabilities based on equations (9) and (10).

Using the equations (9) and (10), the predicted jumping height *h* can be written as

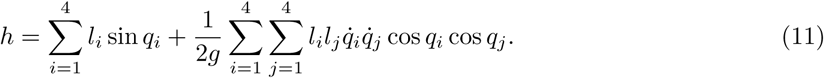

The first candidate input data for the ridge regression were the joint angles and angular velocities (blue line in Fig. 4A). The second candidate data were the functions in the forward kinematics of the position and velocity of the hip joint (equations (9) and (10), red line in Fig. 4A). The third candidate data were the functions that appeared in equation (11) (orange line in Fig. 4A). In GEM (blue crosses in Fig. 4B), we calculated the task-relevant and task-irrelevant variabilities based on equation (11).

### Relation between ridge regression and principal component analysis

It was possible to analytically determine the relation between the ridge regression and principal component analysis (PCA) by decomposing ***X*** using singular value decomposition (SVD), ***X*** = ***UDV*** ^*T*^, where ***U*** ∈ **R**^*T×T*^ is an orthogonal matrix, ***D*** ∈ **R**^*T×D*^ includes the square root of the *i*th eigenvalue of ***X***^*T*^ ***X*** at (*i, i*) element and *D*_*i,j*_ = 0 when *i* ≠ *j*, and ***V*** ∈ **R**^*D×D*^ is an orthogonal matrix. Using the SVD and equation (6), the predicted output *h*_*t*_ can be written as

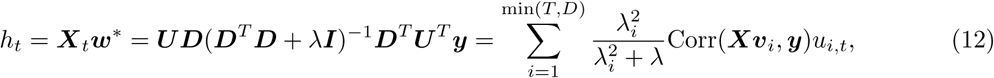

where min(*T, D*) determines the rank of ***X***, 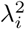 is an eigenvalue of ***X***^*T*^ ***X***, Corr(·, ·) indicates the correlation between two vectors, ***v***_*i*_ is the eigenvector of ***X***^*T*^ ***X*** corresponding to 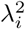, and *u*_*i,t*_ is the (*i, t*) component of ***U***. On the other hand, PCA enables the decomposition of ***X***_*t*_ as

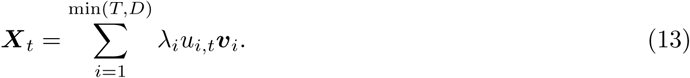

This equation indicates that the motion data can be decomposed into eigenvectors (principal components) with weight *λ*_*i*_*u*_*i,t*_. By comparing equations (12) and (13), the ridge regression enables the prediction of output data by weighting based on the *i*th eigenvector with weight 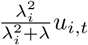 (notably, 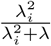 was a monotonic function concerning *λ*_*i*_). An important difference between PCA and the ridge regression is whether the task relevance of the *i*th eigenvector, Corr(***Xv***_*i*_, ***y***), should be considered. Although PCA relies only on the eigenvalue, the ridge regression considers both (nonlinearly transformed) eigenvalue and task relevance. The ridge regression could thus be considered an extended version of PCA to determine how each principal component is relevant to the task.

In PCA, we found the relation between the explained variance and prediction power to be as follows: at a *z*% explained variance, we determine the number of principal components based on 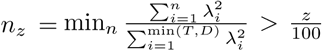 (i.e., the minimum number of principal components that exceed z% explained variance). After determining *n*_*z*_, motion data can be reconstructed as 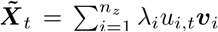. We then multiply 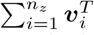 by the 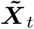 from the right-hand side, resulting in 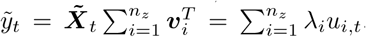 Finally, we calculate the correlation between observed jumping height *y*_*t*_ and 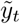 in Figs. 4D-4F.

## Supporting information

## Acknowledgement

This work was supported by a Grant-in-Aid for Young Scientists (18K17894). We thank D. Nozaki and S. Hagio for their helpful comments.

## Data availability

All data generated or analyzed during this study are included in this article.

## Author contributions

D.F. and K.T. designed and performed the experiments. D.F. and K.T. performed the analyses and wrote the manuscript.

