## Supplementary material for "Detection of task-relevant and task-irrelevant motion sequences: application to motor adaptation in goal-directed and whole-body movements": TaskRelIrr.m

```
% Detecting the performance-relevant and performance-irrelevant motion
% sequence
% Furuki & Takiyama, 2018
% ----
% @ Ken Takiyama, Tokyo Univ. Agri & Tech, 2018, 11, 15
% ----

% --- ex. ---
% clear all; close all; clc; 
% load DatDam.mat
% [Xstr, Vstr, ystr, wstr, Estr] = TaskRelIrr(X, y); 
% --- ex. ---

function [Xstr, Vstr, ystr, wstr, Estr] = TaskRelIrr(X, y) 

    % No normalization is necessary in X and y
    % X ... vectorized motion data (the size of X should be T~D)
    % y ... performance data (the size of X should be D~P)
    % T ... # of trials
    % D ... # of dimensions of X
    % P ... # of performance (we recommend that it should be set to 1 [i.e., scalar performance value in each trial])
    % When the number of joints is J and that of time frames is F, D = J~F when you focus on joint angles. 
    % D = 2~J~F when you focus on joint angles and angular velocities. 
    % For example, X(t,:) = (q1(t,:), q2(t,:), ..., qJ(t,:), dq1(t,:), dq2(t,:), ..., dqJ(t,:))

    % Xstr.Xtot, Xstr.Xrel, and Xstr.Xirr are normalized
    % Xstr.Xtot  ... normalized X
    % Xstr.Xrel  ... task-relevant motion sequence
    % Xstr.Xirr  ... task-irrelevant motion sequence
    % Vstr.Vrel  ... task-relevant variability
    % Vstr.Virr  ... task-irrelevant variability
    % Vstr.Cov   ... covariance between task-relevant and task-irrelevant
    % components
    % ystr.ytot  ... normalized y
    % ystr.yTest ... normalized y for test
    % ystr.]yPred ... predicted y
    % Estr.Err_cv ... prediction error in cross validation
    
    % -- Parameters for cross validation -- 
    Itefold    = 10; % # of cross validation
    Ite_cv     = 5;  % # of iterations in cross validation
    lambda_seq = exp(linspace(-7,7,120)); % Regularization parameter for cross validation
    rng('shuffle') % shuffle
    Itefold    = round(Itefold); 
    Ite_cv     = round(Ite_cv); 
    % -- Parameters for cross validation -- 


    [T, D]   = size(X); % size of X
    [Ty, Dy] = size(y); % size of y
    % --- Ridge regression ---
    for ite_loop = 1:Ite_cv
        ar_seq = randperm(T); % for cross validation
        eyeX   = eye(D);      % unit matrix for ridge regression
        for ite = 1:length(lambda_seq)
            lambda = lambda_seq(ite); 

            for ite_cv = 1:Itefold
                seqTest  = ar_seq((floor(T./Itefold)*(ite_cv-1)+1):floor(T./Itefold)*ite_cv); % for test data[
                seqTrain = setdiff(1:T, seqTest); % for training data

                XTest  = X(seqTest, :); 
                XTrain = X(seqTrain, :); 
                yTest  = y(seqTest); 
                yTrain = y(seqTrain); 
                
                % -- size check -- 
                [TTest, DTest]   = size(XTest); % size of X
                [TyTest, DyTest] = size(yTest); % size of y
                if TTest ~= TyTest
                    disp('The number of rows in X and y shoulbe the same'); 
                    return; 
                end
                
                [TTrain, DTrain]   = size(XTrain); % size of X
                [TyTrain, DyTrain] = size(yTrain); % size of y
                if TTrain ~= TyTrain
                    disp('The number of rows in X and y shoulbe the same'); 
                    return; 
                end
                % -- size check -- 

                % -- normalization -- 
                XTest  = (XTest - ones(TTest,1)*mean(XTest,1))./(ones(TTest,1)*std(XTest)); % normalize (0 mean, 1 std)
                yTest  = (yTest - mean(yTest))/std(yTest); % normalize (0 mean, 1 std)
                
                XTrain = (XTrain - ones(TTrain,1)*mean(XTrain,1))./(ones(TTrain,1)*std(XTrain)); % normalize (0 mean, 1 std)
                yTrain = (yTrain - mean(yTrain))/std(yTrain); % normalize (0 mean, 1 std)
                % -- normalization -- 

                w           = inv(XTrain'*XTrain + lambda*eyeX)*XTrain'*yTrain; % weight value for ridge regression
                Err(ite_cv) = (yTest - XTest*w)'*(yTest - XTest*w)./length(seqTest); % prediction error
            end

            Err_cv(ite,ite_loop) = mean(Err); 
        end
    end
    
    % for finding the optimal regularization parameter
    if Ite_cv > 1
        [acv,bcv]      = min(mean(Err_cv,2)); 
    else
        [acv,bcv]      = min(Err_cv); 
    end
    
    % -- normalization -- 
    X = (X - ones(T,1)*mean(X,1))./(ones(T,1)*std(X)); % normalize (0 mean, 1 std)
    y = (y - mean(y))/std(y); % normalize (0 mean, 1 std)
    % -- normalization -- 
    
    lambda_optimal = lambda_seq(bcv); % optimal regularization parameter
    w_opt          = inv(X'*X + lambda_optimal*eyeX)*X'*y; % weight value with optimal lambda
    % --- Ridge regression ---


    % --- Prediction ---
    yPred = []; yTest = []; 
    for ite_cv = 1:Itefold
        seqTest  = ar_seq((floor(T./Itefold)*(ite_cv-1)+1):floor(T./Itefold)*ite_cv); % for test data

        XT  = X(seqTest, :); 
        yT  = y(seqTest); 

        yPred  = [yPred; XT*w_opt]; 
        yTest  = [yTest; yT]; 
    end
    % --- Prediction ---


    % --- data --- 
    Xstr.Xrel   = X*w_opt*w_opt'./norm(w_opt)./norm(w_opt); 
    Xstr.Xirr   = X - Xstr.Xrel; 
    Xstr.Xtot   = X; 
    
    Vstr.Vrel   = mean(var(Xstr.Xrel)); 
    Vstr.Virr   = mean(var(Xstr.Xirr)); 
    Vstr.Cov    = 1 - Vstr.Virr - Vstr.Vrel; 
    
    ystr.ytot   = y; 
    ystr.yPred  = yPred; 
    ystr.yTest  = yTest;
    
    wstr.w_opt  = w_opt; 
    wstr.lambda = lambda_optimal; 
    
    Estr.Err_cv = Err_cv; 
    % --- data --- 

end
```
